# Single-femtosecond atomic-resolution observation of a protein traversing a conical intersection

**DOI:** 10.1101/2020.11.13.382218

**Authors:** A. Hosseinizadeh, N. Breckwoldt, R. Fung, R. Sepehr, M. Schmidt, P. Schwander, R. Santra, A. Ourmazd

## Abstract

The structural dynamics of a molecule are determined by the underlying potential energy landscape. Conical intersections are funnels connecting otherwise separate energy surfaces. Posited almost a century ago ^1^, conical intersections remain the subject of intense scientific investigation ^2–4^. In biology, they play a pivotal role in vision, photosynthesis, and DNA stability ^5,6^. In ultrafast radiationless de-excitation ^1,7^, they are vital to ameliorating photon-induced damage. In chemistry, they tightly couple the normally separable nuclear and electronic degrees of freedom, precluding the Born-Oppenheimer approximation ^8^. In physics, they manifest a Berry phase, giving rise to destructive interference between clockwise and anti-clockwise trajectories around the conical intersection ^9^. Accurate theoretical methods for examining conical intersections are at present limited to small molecules. Experimental investigations are challenged by the required time resolution and sensitivity. Current structure-dynamical understanding of conical intersections is thus limited to simple molecules with around 10 atoms, on timescales of about 100 fs or longer ^10^. Spectroscopy can achieve better time resolution, but provides only indirect structural information. Here, we present single-femtosecond, atomic-resolution movies of a 2,000-atom protein passing through a conical intersection. These movies, extracted from experimental data by geometric machine learning, reveal the dynamical trajectories of de-excitation via a conical intersection, yield the key parameters of the conical intersection controlling the de-excitation process, and elucidate the topography of the electronic potential energy surfaces involved.

---

The quantum mechanical energies of molecular electrons as a function of molecular geometry give rise to effective potential energy surfaces for the motion of the atomic nuclei. When there are *d* nuclear degrees of freedom, the potential energy surface (PES) is *d*-dimensional. In the so-called Born-Oppenheimer (BO) approximation, the electronic and nuclear degrees of freedom are treated separately. In thermally activated chemistry, often only a single BO PES determines the structural dynamics, ensuring the validity of the BO approximation. When two PES’s come into contact, the BO approximation is no longer valid ^11^.

A conical intersection is such a region of such potential energy degeneracy, forming a (*d* − 2)-dimensional manifold with divergent, non-BO coupling between the participating electronic states. The resultant strong mixing of electronic and vibrational degrees of freedom opens a pathway by which dynamical changes of the molecular geometry can cause a transition from one electronic state to another ^11^. As this gives rise to ultrafast, non-radiative relaxation of the excited state, conical intersections play an important role in numerous processes in nature. Trans-to-cis isomerization of the p-cinnamic acid chromophore in Photoactive Yellow Protein (PYP) ^7,12^ (Fig. 1), and retinal ^5,13^ are prime examples.

**Fig. 1.**
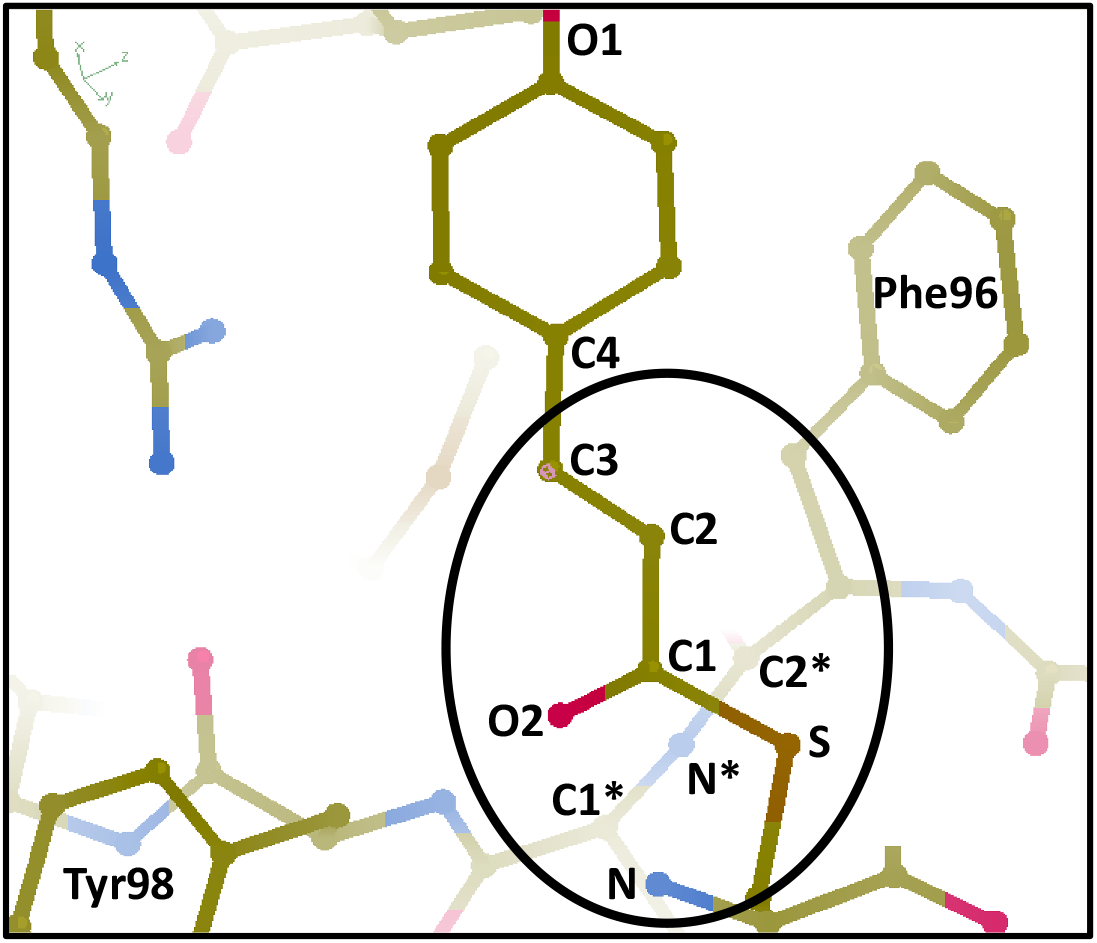
The PYP chromophore and surrounding structures. The oval contains the main structure-dynamically active region, with the numbered atoms and aromatic structures identified. C: carbon, N: nitrogen, O: oxygen, S: sulfur.

Accurate theoretical methods for treating coupled electronic and vibrational dynamics are currently restricted to small molecules. The quality of such simulations – employing either a quantum or a classical treatment of nuclear motions – depends upon the precise characterization and complexity of the PES’s involved. PYP, for example, is composed of 2,289 atoms ^14^, exhibiting 6,861 vibrational degrees of freedom. This level of complexity renders rigorous, first-principles electronic structure calculations unfeasible for the foreseeable future. State-of-the-art density functional theory can be applied to molecules of comparable size ^15^, but does not yet provide reliable chemical accuracy, particularly for conical intersections. Even if it were possible to solve the electronic Schrödinger equation for a single molecular geometry accurately, the total number of molecular geometries needed for adequate sampling of the potential energy landscape as a whole grows exponentially with the number of degrees of freedom.

Experimentally, conclusive observation of the structure-dynamical modes driving electronic switching via conical intersections has remained elusive, because high temporal and spatial resolutions must be combined to resolve the ultrafast dynamics with sufficient acuity. Optical pump-probe spectroscopy provides information on the electronic-state population dynamics with femtosecond time resolution, but it does not offer direct access to the structural properties of the system ^16^. Similarly, time-resolved diffraction techniques, such as ultrafast electron diffraction (UED) and time-resolved X-ray diffraction, have, up to now, lacked the temporal resolution needed to follow de-excitation via a conical intersection ^10,17^. Recently, the combination of UED experiments with extensive, sophisticated ab initio simulations, accomplished the structural characterization of CI-induced dynamics in an 11-atom molecule with a time resolution of 150 fs ^10^.

We report four key advances. First, structure-dynamical collective modes, and trajectories of ultrafast de-excitation can be extracted with atomic spatial resolution and single-femtosecond time resolution from existing time-resolved crystallographic data. For PYP, this information directly reveals that oscillatory charge redistribution on specific molecular bonds surrounding the chromophore plays an important role in the ultrafast de-excitation dynamics of the chromophore.

Second, in combination with tractable and accurate computational methods, the topography of the electronic states involved in de-excitation via a conical intersection can be determined. Third, our approach can be used to determine the key collective variables and boundary conditions controlling the de-excitation dynamics of molecules consisting of thousands of atoms. Finally, the combination of data-driven machine learning with existing experimental and theoretical techniques offers the time resolution of spectroscopy and the spatial resolution of structural methods.

The experimental data were obtained in a time-resolved (optical pump, X-ray probe) serial femtosecond crystallographic study of PYP, as reported in detail elsewhere ^7^. This protein is known to undergo a rapid trans-to-cis isomerization reaction via a conical intersection ^7^. The data consist of a time-series of two-dimensional (2D) diffraction snapshots, each stemming from a different random central slice through the 3D diffraction volume. Each “light” 2D snapshot was recorded after optical excitation at a timepoint known to an accuracy of ~100 fs due to unavoidable “timing jitter” between the optical pump and the X-ray probe pulses ^7,18^. In addition, “dark” snapshots were recorded without any optical excitation. Conventionally, a sufficient number of light 2D snapshots from the same nominal timepoint are indexed and combined (merged) to obtain the 3D diffraction volume and thence the difference between the light and dark atomic structures at each timepoint. (See, e.g., [7].) The timing jitter limits the time resolution of the merged 3D volumes to ~100 fs. This is a severe limitation, because de-excitation via a conical intersection is often complete within that time frame (see below).

We circumvent this problem by applying geometric machine learning ^18–22^ to the same dataset of 2D diffraction snapshots, in order to reconstruct a time-series of 3D diffraction volumes, each pertaining to a timepoint determined with an accuracy of about 1 fs (for details see Methods, and [18]). In essence, our approach rests on the celebrated realization by Takens ^23^ and Packard ^24^ that dynamics tightly constrains the time evolution of a system. This means much less data is needed to reconstruct dynamics than conventionally thought necessary. As an extreme example, Newton’s laws of motion require only one snapshot of the initial conditions (positions and momenta) and the forces acting on a system to predict the dynamical evolution of a non-chaotic system forever. In a similar vein, the time evolution of the diffraction signal is highly constrained by the charge dynamics of the photoexcited system under observation. This allows an essentially jitter-free time-series of 3D diffraction volumes to be recovered from a time-series of 2D central slices, each recorded with substantial timing uncertainty. This algorithmic approach has been validated with experimental data ^18,25,26^, and with data from synthetic models, where the actual “ground-truth” is known ^18,25,26^. (See also Methods.)

Armed with a series of accurately time-stamped 3D diffraction volumes, standard time-resolved crystallographic approaches ^27,28^ can be used to compile jitter-free difference electron density movies, revealing the dynamics of the photo-excited charge distribution. As described in detail in ^18^, using time-lagged embedding ^23,24^, our data-analytical pipeline “learns” the Riemannian manifold on which the dynamics unfolds, and conducts all analysis, including (nonlinear) singular value decomposition, on that curved manifold ^18,21^. This approach yields the characteristic collective modes of the charge distribution (“topos”), and their respective time evolutions (“chronos”). More specifically, each topo represents a characteristic difference electron density map, evolving in time as prescribed by its corresponding chrono (Fig. 2). Each topo-chrono pair thus represents a characteristic dynamical mode of the charge distribution. In essence, these modes constitute the basis functions, which may be used in combination to describe the dynamical trajectories (“the reaction paths”) of the system (see below).

**Fig. 2 | a.**
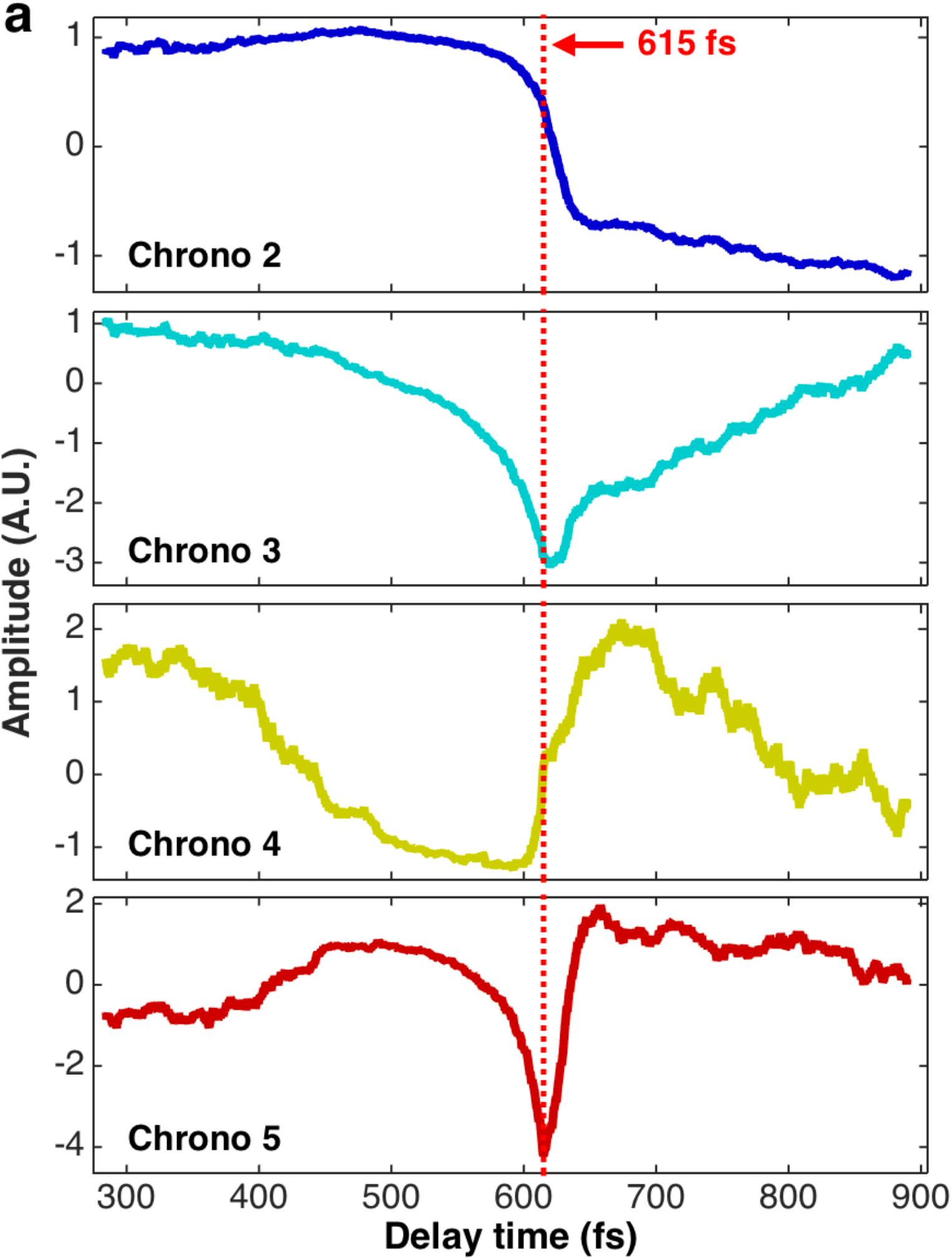
Evolution of dynamical modes (chronos) 2 – 5 as a function of pump-probe delay. Chrono 1 represents the moving average, and is not shown.

Fourier analysis of the chronos by multi-taper methods ^29,30^ reveals the clear presence of frequencies of up to 95 THz (10.5 fs) at signal-to-noise ratios of ~ 8 or more (Supplementary Fig. S5). As the observation of a frequency component in an initially non-uniform set of timepoints requires a time resolution ~5x shorter than the period of the component ^31^, the observation of a clear 10.5 fs signal validates the high time resolution of our approach.

All chronos display a sharp turning point at 615 fs (Fig. 2a). Chronos 2 and 4 display approximate inversion symmetry, while chronos 3 and 5 show approximate mirror symmetry about this timepoint. The corresponding characteristic charge distributions, more precisely the characteristic difference electron density maps are shown in Fig 2b. Movies of the structure-dynamical modes with femtosecond time resolution are shown in Supplementary Movies S1-S4. The sharp (~3-fs-wide) turning points in the chronos (Fig 2a) further corroborate the high temporal resolution of our technique.

**Fig. 2b.**
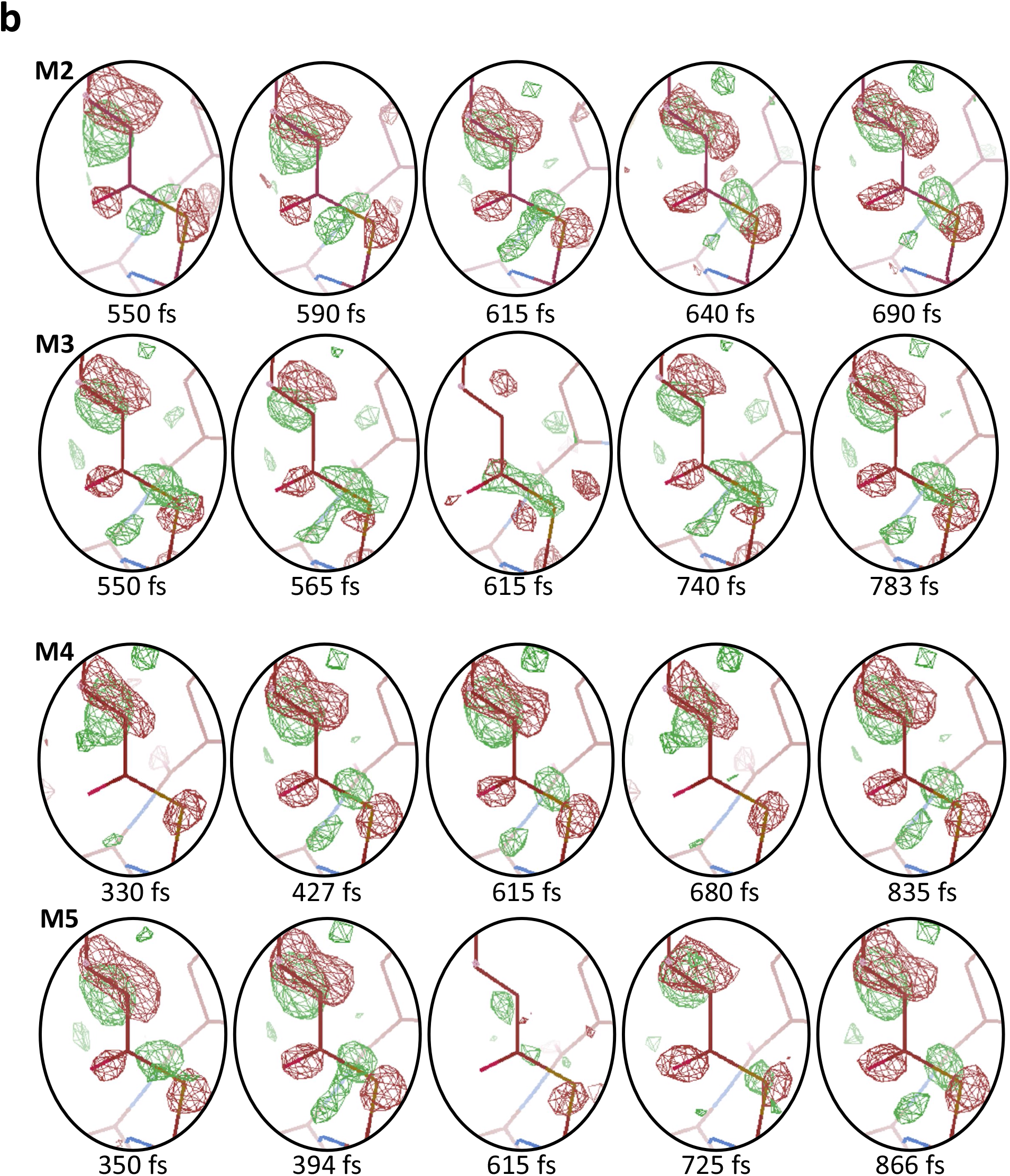
Evolution of difference electron density (DED) with time for dynamical modes M2 – M5. The difference electron density maps near the top of the oval region have previously been associated with trans-to-cis isomerization. The patterns near the bottom of the oval region show strong oscillatory charge dynamics.

We now describe the structure-dynamical information provided by these modes. As Mode 1 represents the moving average of all modes, its topo is added to each of the subsequent modes so as to highlight the deviations from a moving average. We emphasize that the Supplementary Movies 1 – 4 offer the best means of examining the rich variety of dynamical features, which are difficult to illustrate by a small number of individual snapshots.

Mode 2, with a time evolution (chrono) resembling a step function (Fig. 2), represents the only structure-dynamical evolution not reversed within the picosecond timespan of the dataset. The other chronos return roughly to their initial values in the course of the experimental timespan. In Mode 2, the topo represents strong difference electron density features in the vicinity of the C2-C3 and C3-C4 bonds in PYP (Fig. 1), previously associated with the trans-to-cis isomerization ^32^. Importantly, Mode 2 also reveals previously unreported rapid charge oscillations along the bonds between C1*, N*, and C2* connecting the Phe96 and Tyr98 aromatic structures.

The V-shaped chrono of Mode 3 reaches its most negative value at 615 fs. This evolution is reversed in sequence and magnitude over the subsequent 300 fs. In addition to well-known changes in the electron density normally assigned to isomerization, Mode 3 reveals substantial oscillatory charge redistribution along the S – C1 bond in the chromophore, and the bonds connecting C1*, N*, and C2* surrounding the chromophore (Fig. 1).

Mode 4 recapitulates the signature of the difference electron density associated with isomerization, but also shows significant oscillatory charge transfer along the conduit connecting the Phe96 and Tyr98 structures. In contrast, the sign-change of chrono 4 at 615 fs results in the *same* sequence of events after 615 fs as before that timepoint, rather than the reverse sequence of events observed after the sharp turning point in Mode 3.

Finally, Mode 5 displays the changes associated with isomerization and strong oscillatory charge redistribution along the conduit between Phe96 and Tye98. Remarkably, however, the difference electron density map at 615 fs is almost featureless, indicating the structure at that timepoint closely approximates that of the *dark* structure. Taken together, the modes described above reveal the structure-dynamical motifs involved in PYP de-excitation.

As noted earlier, although the characteristic structure-dynamical modes (basis functions) described above display key features of the system, they must be appropriately combined to reconstruct the trajectories (reaction pathways) associated with de-excitation via the conical intersection. Additional information is needed to accomplish this task. As a minimal ansatz, we assume that, in the close proximity of the conical intersection, a trajectory plays out on a two-dimensional PES. The admissible dynamical trajectories can then be determined as follows. We compare the six possible experimental trajectories obtained from pairwise combinations of experimental modes, with trajectories contained in a dictionary of half a million theoretically calculated de-excitation trajectories. The calculations employ an effective model Hamiltonian in a compressed collective-mode space to capture the properties of a conical intersection and its vicinity. This allows numerically exact calculation of the nuclear quantum-dynamics as a function of a small number of molecule-specific model parameters. By comparing the resulting simulated structure-dynamical trajectories with the experimentally determined dynamical trajectories revealed by our data-analytical pipeline, we determine the numerical values that the model parameters assume in PYP, thus gaining detailed insight into the topography of this photochemically exemplary conical intersection (see Methods).

Our theoretical model is tantamount to the simplest possible realization of a conical intersection, implicitly assuming dissipation into additional modes is negligible on the time scale of 100 fs. The PES’s are approximated by a second-order Taylor expansion, known as the vibronic coupling model ^33,34^, and are assumed to be symmetric with respect to both normal modes, possibly differing in their respective vibrational frequencies. Inter-state coupling is mediated via one mode only.

To determine the dynamics in the space spanned by two modes, we numerically solve the time-dependent Schrödinger equation for a wave packet initially occupying the excited electronic state. We take into account a total of six model parameters, including the reference ground state frequency related to the kinetic energy of the wave packet, the respective vibrational frequencies of both potential energy surfaces, the coupling strength, and the initial position of the wave packet relative to the conical intersection. For 500,000 different combinations of these six model parameters, we perform computer simulations, and determine, as a function of time, the expectation values of the position operators associated with each mode. In this way, we obtain dynamical trajectories in the 2D space spanned by two collective modes, denoted *x* and *y* for short.

A dynamical trajectory in this 2D space consists of a time-ordered sequence of points given by (*x*(t), *y*(t)). Each of *x*(t) and *y*(t) is associated with a characteristic difference electron density map (a topo), whose time-varying contribution is determined by the associated chrono. Thus, in the space spanned by *x* and *y*, a potential dynamical trajectory is obtained by plotting two chronos against each other. Of course, the choice of characteristic modes is not unique; any linear combination, with or without sign inversion and scaling, provides an equally good basis set.

Using the bank of 500,000 simulated de-excitation trajectories in the vicinity of a conical intersection, we identify the experimental trajectory leading to high-probability de-excitation of PYP via a conical intersection. Specifically, we fit each of the simulated trajectories to the six experimental trajectories, using the smallest χ^2^ to identify the best match in each case (Fig. 3a, and Methods). In this, we consider both temporal and spatial translations of the simulated trajectories, and allow the simulated trajectories to be linearly scaled. Following this procedure, we are able to reproduce the observed collective-mode behavior, determine the geometric properties of the conical intersection, and the uncertainties in our determination of these physically important parameters (Table 1 and Methods).

**Table 1.** Parameters of the manifold of potential energy surfaces extracted from the observed five de-excitation trajectories in the neighborhood of the conical intersection. Legend: (r_0_, θ_0_): distance and angle from starting point to the conical intersection; λ: coupling strength; ω: frequency (kinetic energy); γ_1,2_: frequency on respective PES. The superscript *Q* indicates the accuracy is limited by the spacing in the parametric grid of simulated trajectories.

| Trajectory<br>Parameter | M2 – M3 | M2 – M4 | M2 – M5 | M3 – M4 | M4 – M5 |
| --- | --- | --- | --- | --- | --- |
| $r_0$ | $4.11 \pm 2.12$ | $5.22 \pm 1.40$ | $4.67 \pm 1.24$ | $3.56 \pm 0.98$ | $3.00 \pm 1.67$ |
| $\theta_0$ | $-140 \pm 18$ | $-100 \pm 26$ | $-120 \pm 20^{(Q)}$ | $-120 \pm 6.7$ | $-120 \pm 20^{(Q)}$ |
| $\lambda$ | $1.5 \pm 0.27$ | $1.0 \pm 0.33$ | $1.5 \pm 0.5^{(Q)}$ | $1.00 \pm 0.5^{(Q)}$ | $1.5 \pm 0.5^{(Q)}$ |
| $\omega [\times 1E3]$ | $2.50 \pm 0.93$ | $1.67 \pm 1.06$ | $2.08 \pm 0.42^{(Q)}$ | $3.33 \pm 0.28$ | $2.08 \pm 0.42^{(Q)}$ |
| $\gamma_1$ | $0.13 \pm 0.03$ | $0.08 \pm 0.04$ | $0.08 \pm 0.01$ | $0.10 \pm 0.02$ | $0.07 \pm 0.02$ |
| $\gamma_2$ | $0.10 \pm 0.02$ | $0.20 \pm 0.10$ | $0.12 \pm 0.02$ | $0.08 \pm 0.04$ | $0.15 \pm 0.03$ |

**Fig. 3 a - d.**
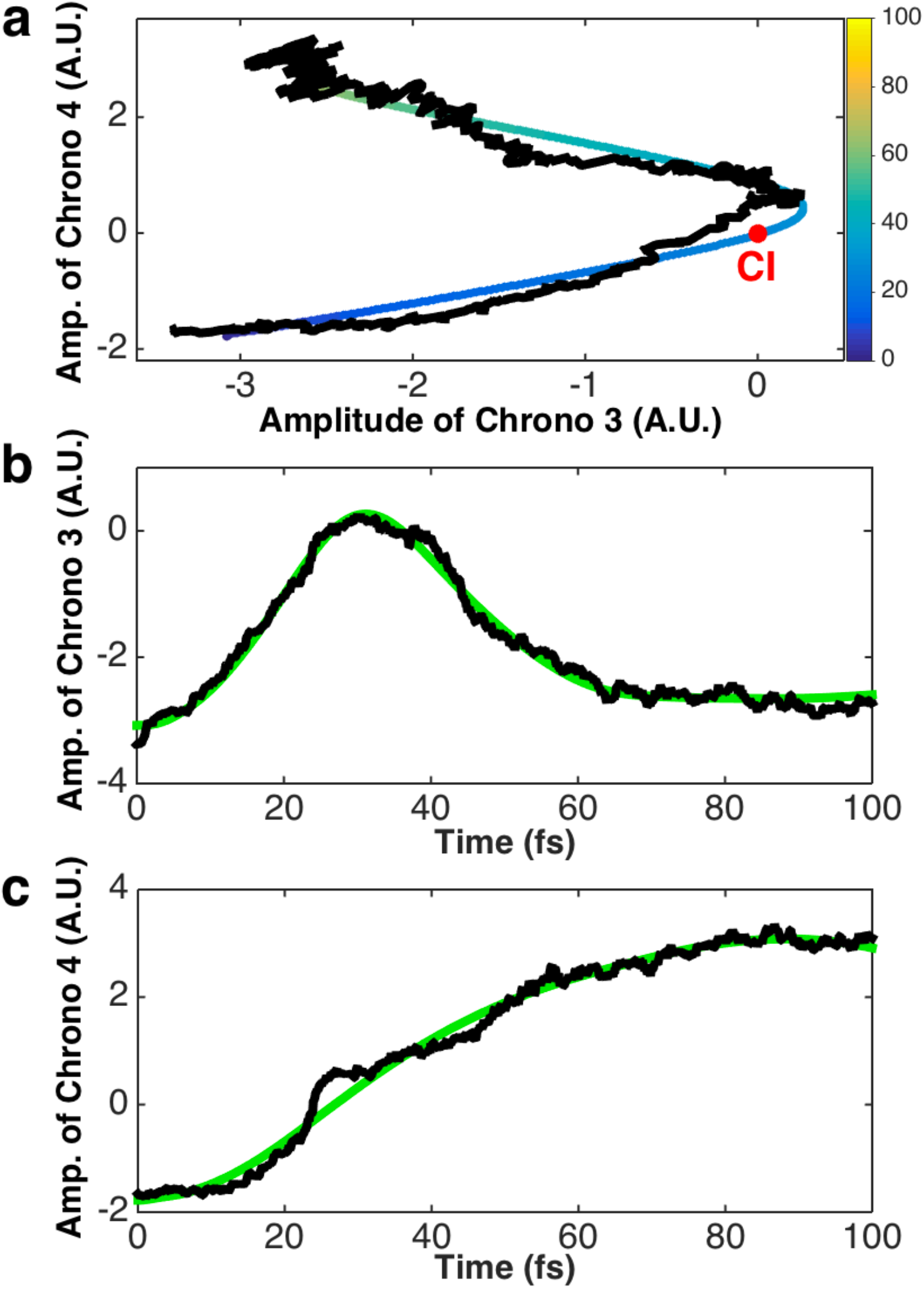
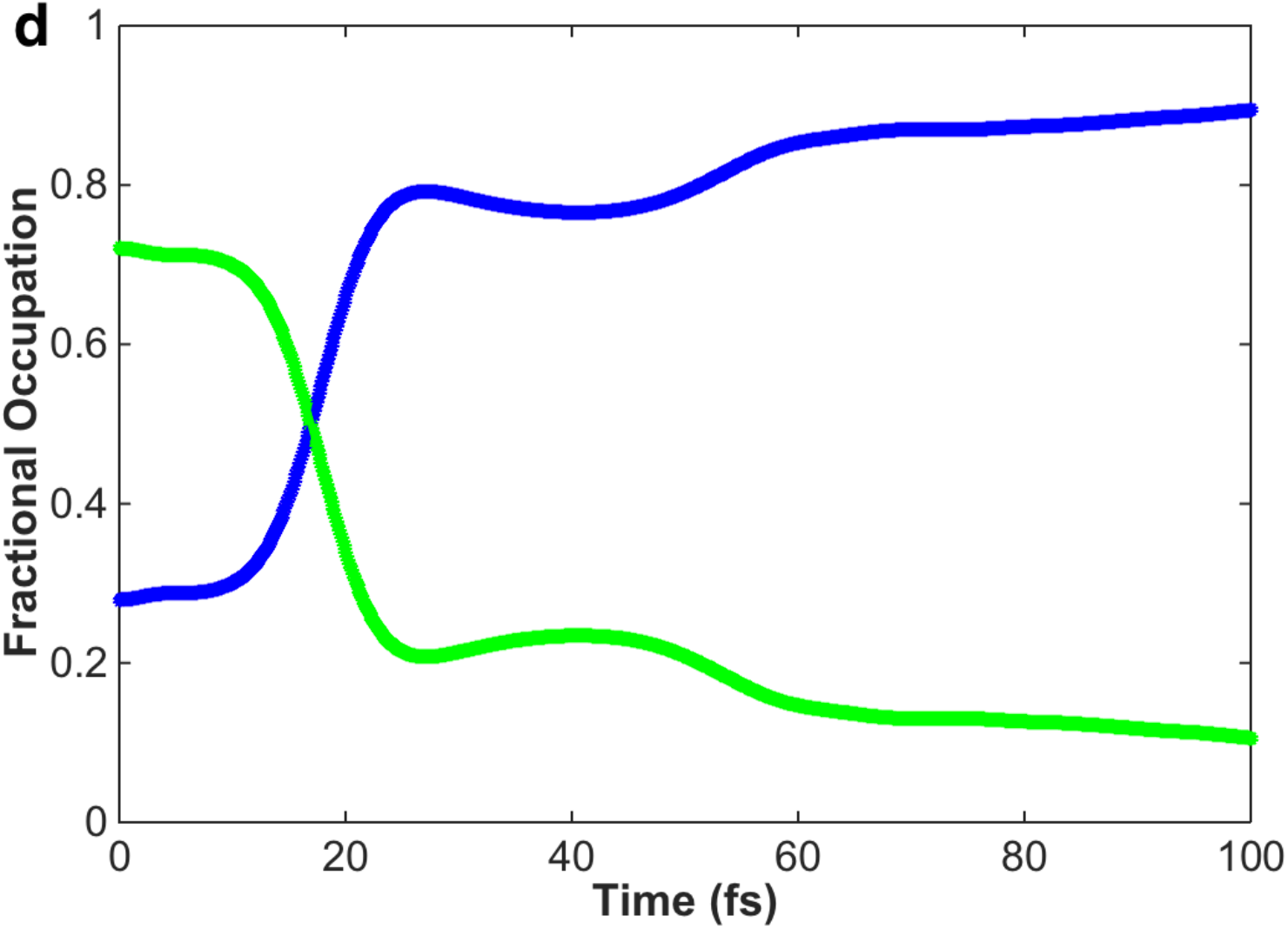
Dynamical trajectories near the conical intersection. **a,** An experimental dynamical trajectory obtained from modes 3 and 4 (in black), and the best-fit simulated trajectory, with color bar showing passage of time. **b, c,** Time evolution along collective modes *x* and *y*, respectively. For additional trajectories, see Supplementary Information. **d,** The calculated de-excitation dynamics as reflected in the electronic state population vs. time for the trajectory of Fig.3a. The green and blue curves represent the populations of the upper and the lower adiabatic electronic states, respectively.

We now discuss results of these fits. Of the six possible pairwise combinations of experimentally determined dynamical modes, five can be identified with simulated trajectories of de-excitation via the conical intersection (Fig. 3a, and Supplementary Figures 6 - 9). (The sixth combination does not correspond to any simulated trajectory in our databank.) The identification of the experimental trajectories with their simulated counterparts yields the key parameters governing the properties of the experimentally investigated conical intersection (Table 1). These parameters provide insight into de-excitation dynamics (Fig. 3b) and the topography of the conical intersection in PYP (Fig. 4). The very similar parameters obtained from all five pairwise combinations of experimentally determined structure-dynamical modes indicate that in the vicinity of the conical intersection, the five dynamical trajectories can be described in terms of the same underlying conical intersection. Further away from the conical intersection, however, our analysis reveals the presence of at least five distinct trajectory segments. We interpret these segments as pertaining to different conduits to and from the vicinity of conical intersection. In the vicinity of the conical intersection, the trajectories represent high-probability de-excitation routes on the same two-dimensional PES manifold.

**Fig. 4.**
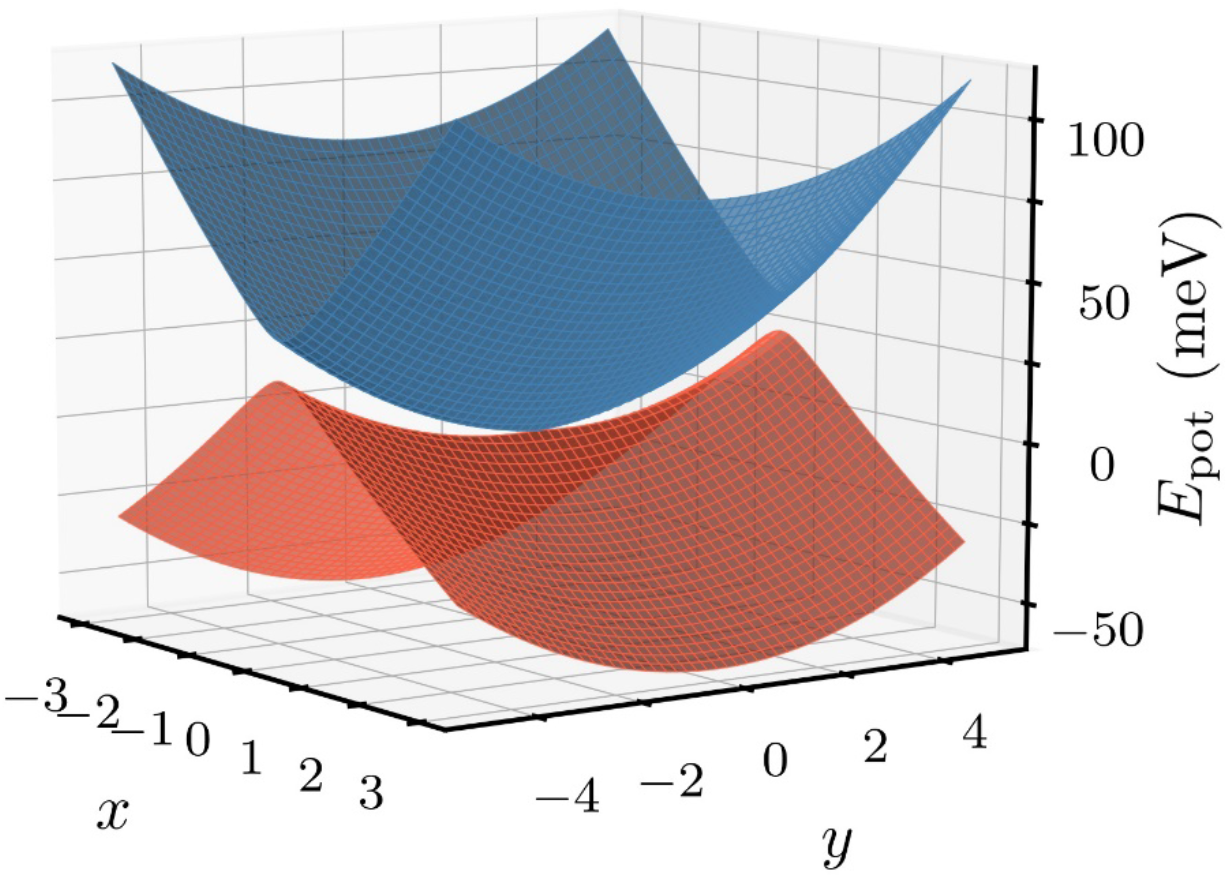
Topography of the conical intersection in PYP, as deduced from five dynamical trajectories. The parametric values deduced from the five experimental dynamical trajectories identified in this paper are shown in table 1. The coordinates, *x* and *y*, representing the normal modes of the employed vibronic coupling model, are shown in dimensionless coordinates.

We now summarize the primary conclusions of our work. First, our results demonstrate a novel data-driven approach, which combines the superb spatial resolution of structural methods, such as crystallography, with the exquisite time resolution of spectroscopy. In essence, this approach is tantamount to structure-dynamical spectroscopy with atomic spatial resolution and femtosecond timing acuity. Second, our results on the atomic-level changes associated with the femtosecond de-excitation of a protein via a conical intersection reveal significant, previously unobserved oscillatory charge dynamics involving bonds to the aromatic structures surrounding the chromophore. This demonstrates that new information can be extracted from existing data with unprecedented spatial and temporal resolution. Finally, by combining geometric machine learning analysis of experimental data with simple and numerically accurate quantum-dynamics simulations, we have demonstrated a powerful new route to studying a wide variety of important processes in complex molecular systems inaccessible by first-principles calculations.

Of course, future tasks remain. These include investigating the possible effect of crystallinity on the observed relaxation modes and trajectories, and whether the small number of important collective variables revealed by our approach offers a route to more accurate theoretical calculations than hitherto possible. These future tasks notwithstanding, our present results already reveal the unanticipated trove of information, which can be extracted from existing experimental data by a combination of data-driven machine learning and physically-based theory.

## ACKNOWLEDGMENTS

We acknowledge valuable discussions with E. Lattman, K. Moffat, T. Martinez, and J.C.H. Spence, and past and present members of the UWM data analysis group. The development of underlying techniques was supported by the US Department of Energy, Office of Science, Basic Energy Sciences under award DE-SC0002164 (underlying dynamical techniques), and by the US National Science Foundation under awards STC 1231306 (underlying data analytical techniques) and 1551489 (underlying analytical models). NB and RS were supported by the Cluster of Excellence ‘CUI: Advanced Imaging of Matter’ of the Deutsche Forschungsgemeinschaft (DFG) – EXC 2056 – project ID 390715994.

## AUTHOR CONTRIBUTIONS

AO designed this study. AO, AH, PS, RF, and RS co-defined the algorithm and the data-analytical pipeline. AH, and RF performed analytical and computational work, and tested and validated the algorithm with participation by AO. NB and RS performed the quantum-dynamics simulations. MS provided experimental data, and expertise in crystallographic data analysis. All authors contributed to the manuscript.

## AUTHOR INFORMATION

The authors declare no financial interests.

## DATA AVAILABILITY

The data are available at: CXI-DB ID 118.

## CODE AVAILABILITY

The code will be made available to the general public upon acceptance of this paper for publication.

## Supplementary Information

**Supplementary Figs. S1 – S10 |** Supplementary figures

**Supplementary Movies S1 – S5 |** Single-mode difference electron density movies

**Supplementary Movies S6 – S9** | Trajectory difference electron density movies

